## Supplemental Figures for "Short heat shock factor A2 confers heat sensitivity in *Arabidopsis*: Insights into heat resistance and growth balance"

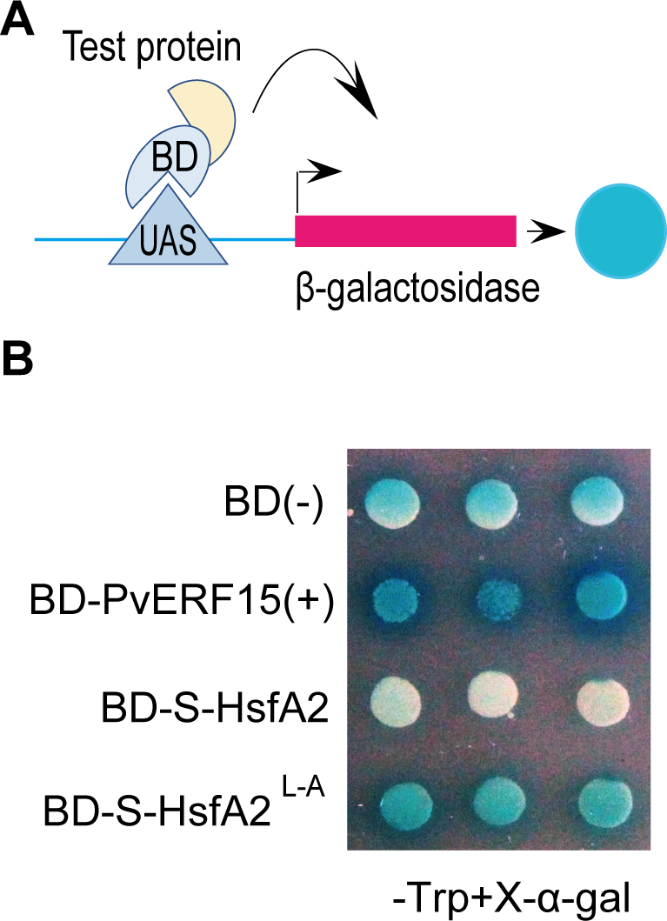

**Figure S1. The** **LxLxLx motif is responsible for S-HsfA2 transcriptional repression.**

**(A)** The working model for the transcriptional activation activity of test proteins in yeast cells, in which the test protein-GAL4 transcription factor DNA binding domain (BD) fusion protein could mediate the UAS element to activate the expression of the *β-galactosidase* reporter gene, eventually generating blue yeast cell clones. **(B)** A yeast strain with the *β-galactosidase* gene driven by a UAS-containing promoter was transformed with a plasmid encoding the GAL4 BD without (BD, negative control) or with a protein (PvERF15 positive control, S-HsfA2, or S-HsfA2^L-A^) fusion. Transformants were subsequently grown on Trp-deficient media supplemented with 5-bromo-4-chloro-3-indolyl-α-D-galactopyranoside (X-α-gal).

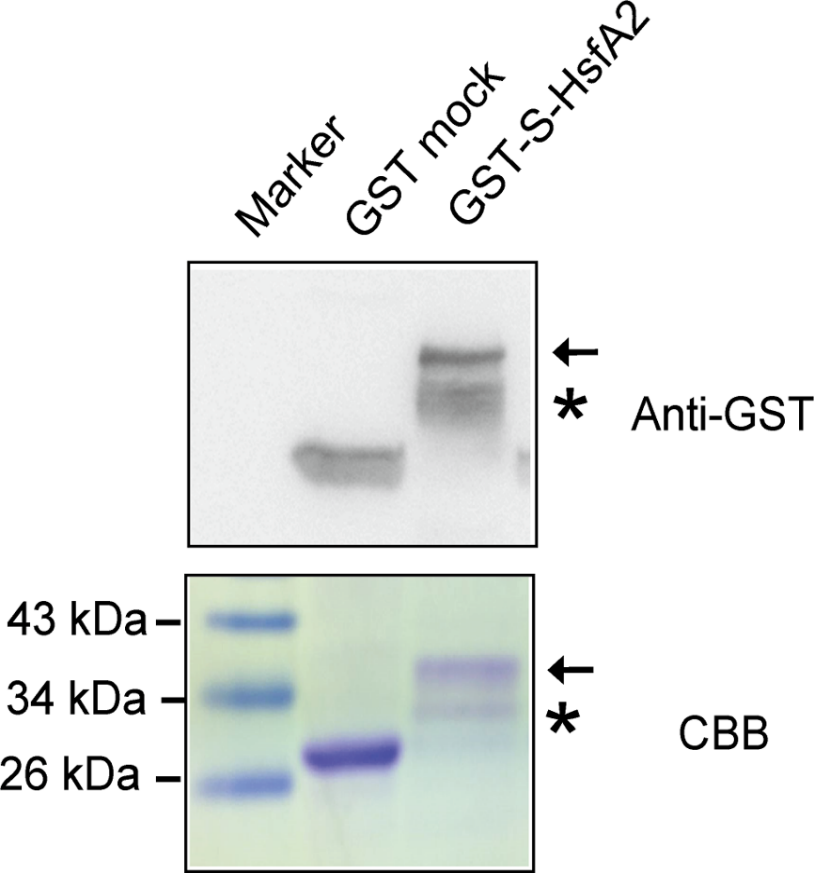

**Figure S2. Bacterially expressed and purified GST-S-HsfA2.**

GST-S-HsfA2 production was confirmed by immunoblotting with an anti-GST antibody. The mock proteins (empty vector control) were used as a negative control. Coomassie Brilliant Blue (CBB)-stained proteins are shown as a loading control. The GST-S-HsfA2 protein signal is indicated by an arrow. The asterisk indicates a nonspecific signal.

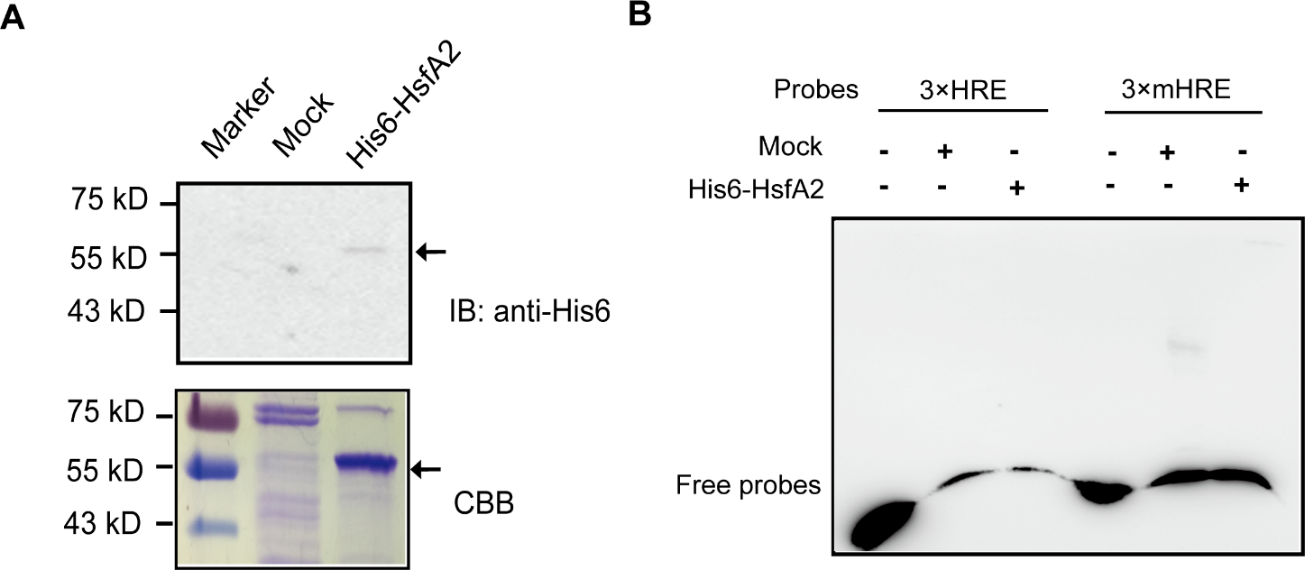

**Figure S3. His6-HsfA2 fails to bind to the HRE *in vitro*.**

**(A)** The production of His6-HsfA2 was confirmed by immunoblotting with an anti-His6 antibody. The mock proteins (empty vector control) were used as a negative control. Coomassie Brilliant Blue (CBB)-stained proteins are shown as a loading control. **(B)** His6-HsfA2 binding to HRE or mHRE was verified by EMSA.

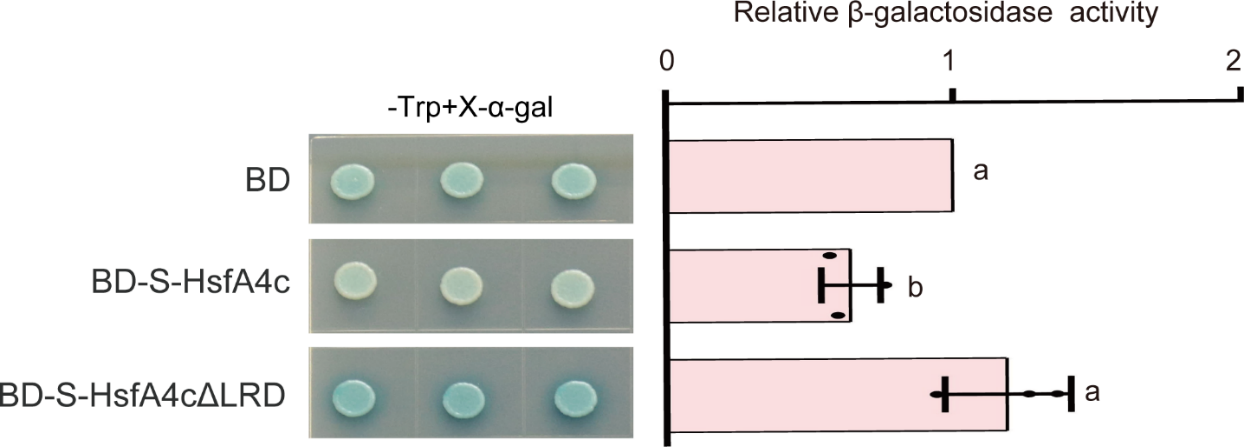

**Figure S4. The** **LRD is responsible for S-HsfA4c transcriptional repression.**

A yeast strain with the *β-galactosidase* gene driven by a GAL4 binding UAS-containing promoter was transformed with a plasmid encoding the GAL4 BD without (BD, negative control) or with a protein [S-HsfA4c or S-HsfA4c lacking LRD (S-HsfA4c△LRD)] fusion. Transformants were subsequently grown on Trp-deficient media supplemented with 5-bromo-4-chloro-3-indolyl-α-D-galactopyranoside (X-α-gal). The β-galactosidase activity was expressed as a ratio relative to the BD, which was set to a value of 1. The data are presented as the means ± SDs of three independent clones. Different letters indicate significant differences according to one-way ANOVA, *P* < 0.05.

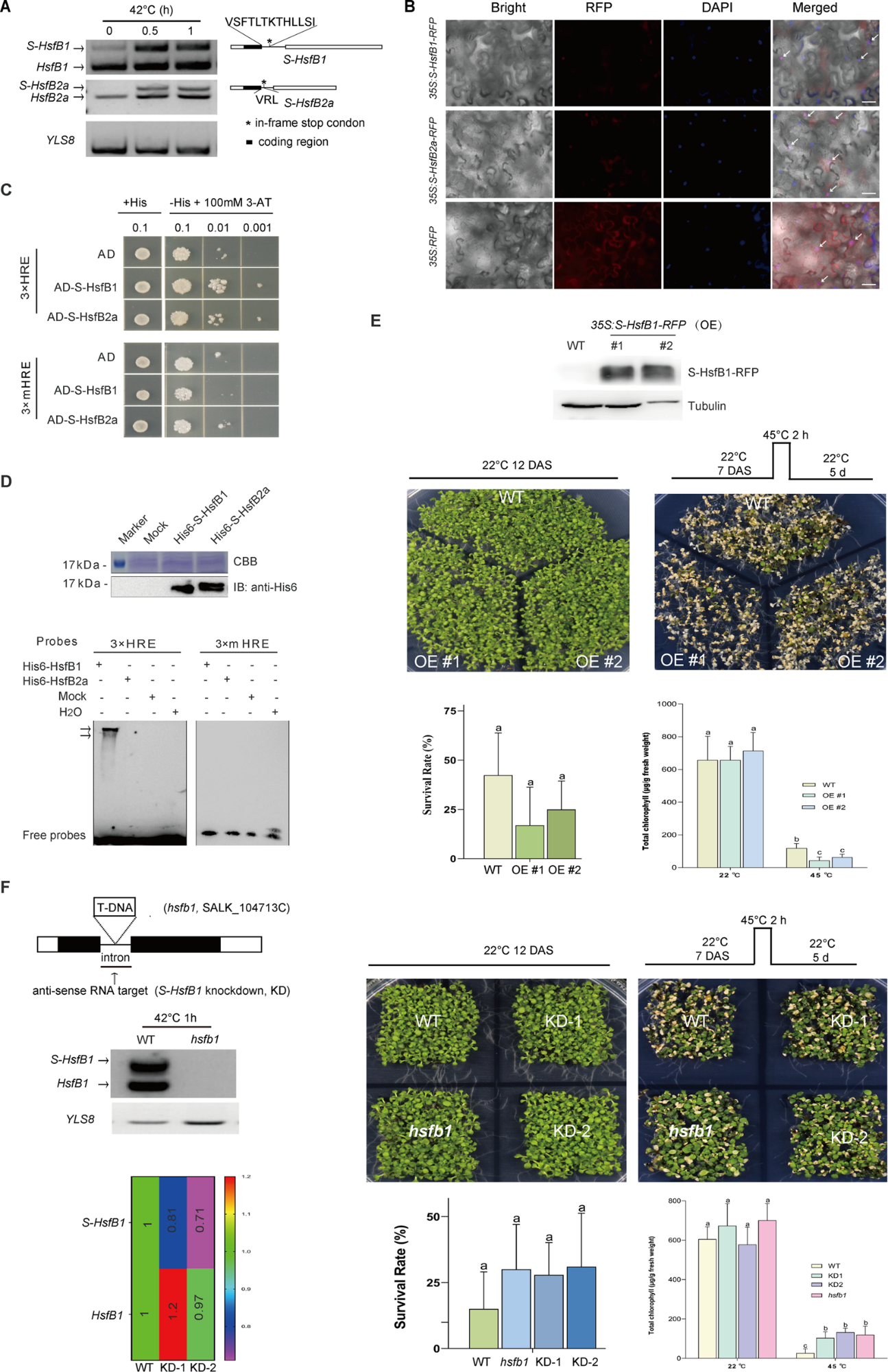

**Figure S5. S-HsfB1 is similar to S-HsfA2 and represents a new HSF.**

**(A)** RT‒PCR splicing analyses of *HsfB1* and *HsfB2a* in 7-d-old *Arabidopsis* plants under heat stress. The *YLS8* gene served as a loading control. The structures of the S-HsfB1 and S-HsfB2a splice variants are shown in the right panel. The amino acid residues encoded by the retained introns are indicated above each variant. **(B)** Representative images showing the subcellular localization of S-HsfB1-RFP or S-HsfB2a-RFP in *N. benthamiana* epidermal cells. RFP was used as a negative control. DAPI, 4,6-diamidino-2-phenylindole (DAPI; nuclei staining). Scale bar, 50 μm. **(C, D)** S-HsfB1 or S-HsfB2a binding to the HRE was verified by Y1H and EMSA with bacterially expressed and purified His6-S-HsfB1 or His6-S-HsfB2a. Mock, empty vector control. **(E)** The *35S:S-HsfB1-RFP*-overexpressing (OE) lines were confirmed by western blot analysis and then subjected to thermotolerance assays. **(F)** The T-DNA *HsfB1*-knockout mutant (*hsfb1-1*) confirmed by RT‒PCR and the antisense (targeting the retained intron sequences)-mediated *S-HsfB1*-knockdown lines (KD-1 and KD-2) verified by RT‒qPCR were subjected to thermotolerance assays. The data are presented as the means ± SDs of three independent experiments. Different letters indicate significant differences according to one-way ANOVA, P < 0.05. DAS, days after sowing.

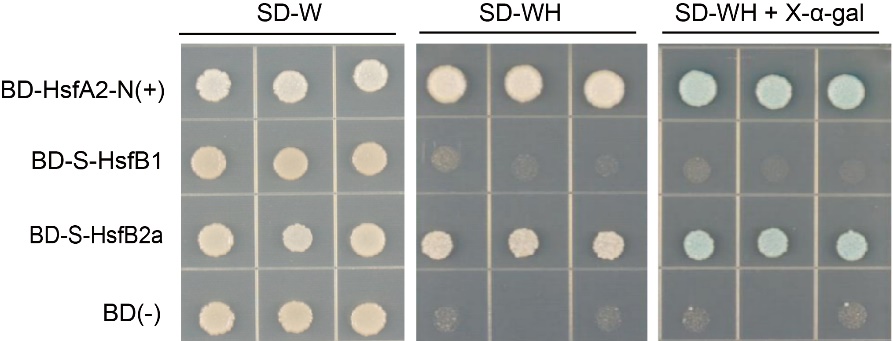

**Figure S6. Transactivation activity of S-HsfB1 and S-HsfB2 in yeast cells.**

A yeast strain with the *β-galactosidase* gene driven by a UAS-containing promoter was transformed with a plasmid encoding the GAL4 BD without (BD, negative control) or with a protein (	HsfA2-N positive control, S-HsfB1, or S-HsfB2a) fusion. Transformants were grown on Trp-deficient or His-deficient media supplemented with 5-bromo-4-chloro-3-indolyl-α-D-galactopyranoside (X-α-gal).

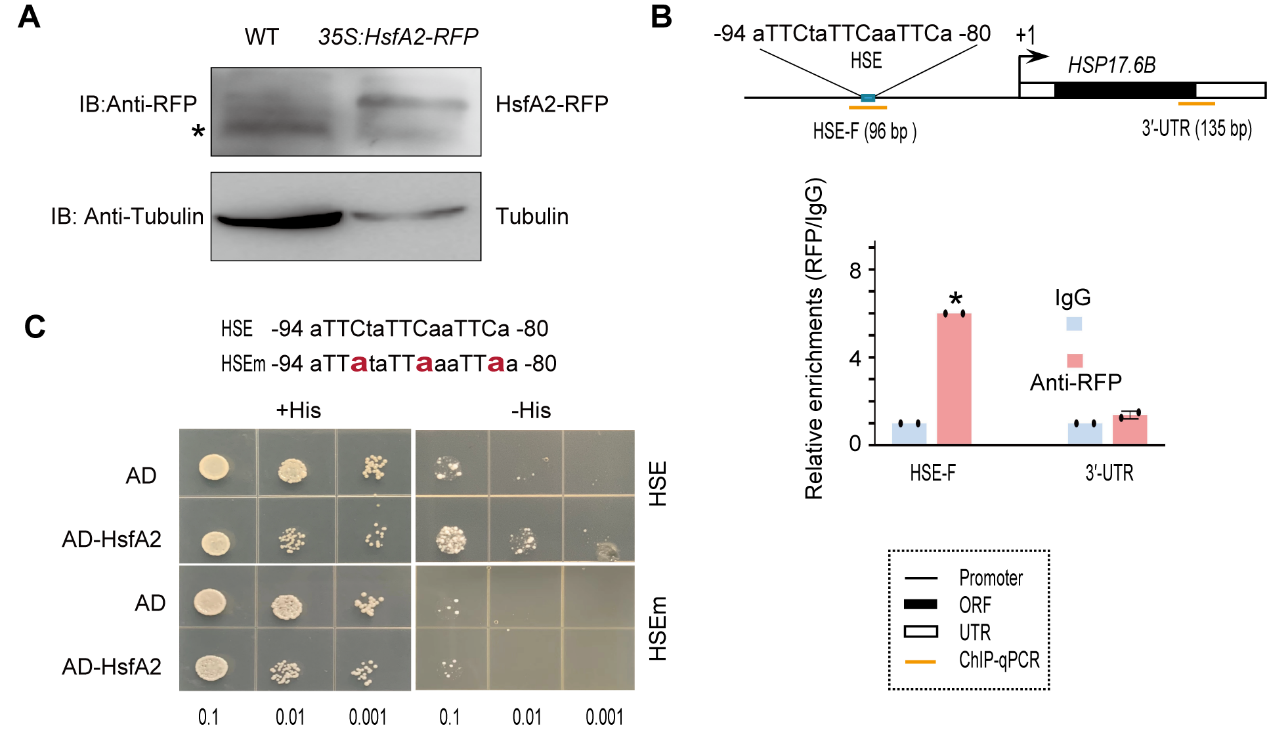

**Figure S7. HsfA2 binds to the HSE of the *HSP17.6B* promoter.**

**(A)** Western blot analysis verified the transient expression of the RFP-tagged HsfA2 fusion protein under the control of 35S (*35S:HsfA2-RFP*). The asterisk indicates a nonspecific signal. **(B)** ChIP experiments were performed with an RFP antibody and mouse IgG (mock control) on Arabidopsis seedlings expressing *35S:HsfA2-RFP*. A diagram of *HSP17.6B* is shown at the top. Relative enrichment was calculated by comparing GFP antibody-immunoprecipitated DNA with that immunoprecipitated with the IgG control, in which IgG was set to a value of 1. The data are presented as the means ± SDs of at least two independent qPCR experiments. The significance of differences between the experimental values was assessed by Student’s *t* test (**P* <0.05 and ***P* <0.01). **(C)** HsfA2 binding to the HSE within HSP17.6Bp was verified by a Y1H assay. The mutated HSE (mHSE) was used as a negative control.

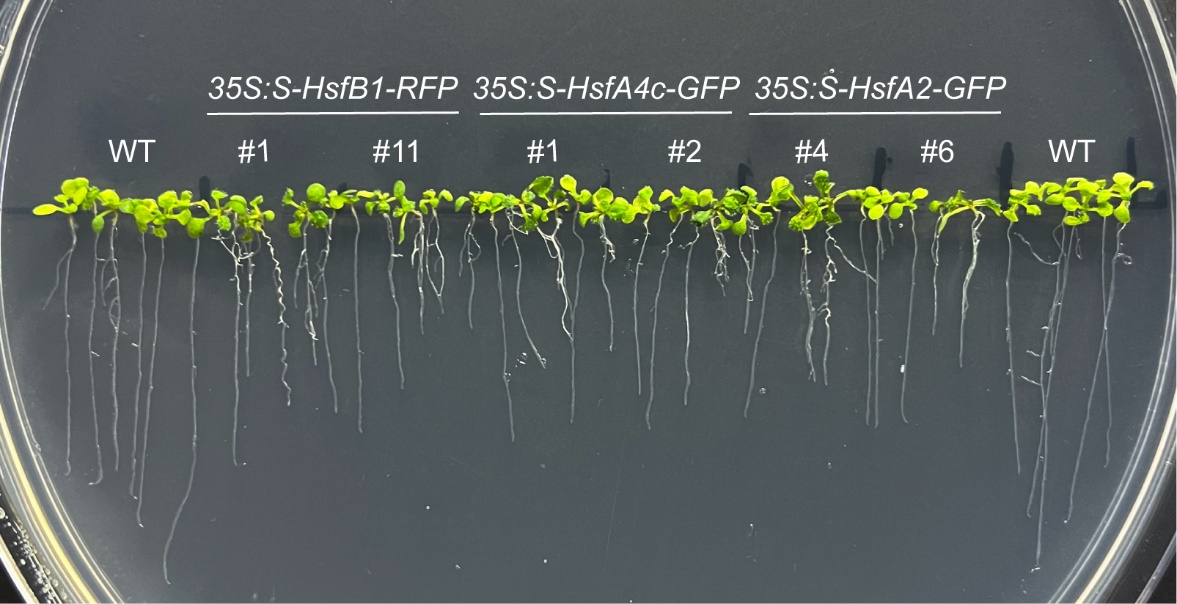

**Figure S8.** **Overexpression of S-HsfA2-GFP, S-HsfA4c-GFP, or S-HsfB1-RFP inhibits the transgenic *Arabidopsis* root growth.**

The seeds of wild-type (WT) control, *35S:S-HsfA2-GFP* lines, *35S:S-HsfA4c-GFP* lines, or *35S:S-HsfB1-RFP* lines were sowed in 1/2 MS media for 14 days. A representative image of root length is shown.
